# Effects of Ankle Stiffness on Total Leg Kinematics, Mechanics, and Muscle Activation during Walking

**DOI:** 10.64898/2026.06.11.731479

**Authors:** Rachel Gehlhar Humann, Michael J. Rose, Will Flanagan, Laila Harris, Alyssa Tomkinson, Alexandra S. Voloshina, Tyler R. Clites

## Abstract

**Purpose:** Ankle stiffness can be altered by normal aging, bone and joint pathology, and treatments such as orthoses or surgical joint fusion. The effects of ankle stiffness on gait are not yet well understood but may be crucial for understanding how these pathologies and treatments influence body mechanics. The objective of this work was to investigate how isolated changes in stiffness applied in parallel with the ankle impact lower-limb kinematics, kinetics, joint work, and muscle activation during walking in individuals without lower-limb pathology.

**Methods:** Nine young adults without lower-limb pathology wore an adjustable-stiffness ankle exoskeleton and walked at 31 different conditions of ankle spring stiffness, neutral angle, and treadmill incline. We recorded motion capture data, ground reaction forces, and muscle activation, and analyzed the resultant data for trends as a function of ankle stiffness.

**Results:** Exoskeleton-side ankle range of motion decreased and asymmetry increased across all joints as ankle stiffness increased, primarily due to decreased plantarflexion at toe-off. The 30 Nm/rad spring stiffness condition led to a minimum in mean exoskeleton-side muscle activation and hip joint work, but increased kinematic asymmetry.

**Conclusion:** Our results suggest that there may exist a range of stiffnesses at the lower end of typically-studied values that can reduce muscle activation and joint work during walking, though at the cost of kinematic symmetry. These findings provide a deeper understanding of how ankle stiffness influences gait mechanics, with potential applications in wearable devices, clinical rehabilitation, and assistive technology.

## 1 Introduction

Changes in ankle stiffness can arise due to certain pathologies, such as arthritis, ligament rupture, and spasticity which disrupt innate tissue compliance, thereby altering normal joint mechanics [1]. For example, changes in ankle joint stiffness have been observed in individuals with stroke-related hemiparesis [1–3], spinal cord injury [4], and rheumatoid arthritis [5]. In addition, clinical interventions used to assist impaired joint function post injury, such as orthoses [6–10] or joint fusion [11, 12], can further modulate joint stiffness in order to compensate for neuromuscular deficits that may impair active torque production.

Altered ankle stiffness has critical consequences for whole-body locomotor mechanics. Evidence from studies of passive devices, including passive ankle exoskeletons and ankle-foot orthoses (AFOs), demonstrates that modifying ankle stiffness can reduce the metabolic cost of walking, increase walking speed, reduce perceived fatigue, and improve user satisfaction [6, 13–15]. However, these benefits strongly depend on how well device stiffness is matched to the individual user, including factors such as body weight, walking speed, activity level, and severity of muscle weakness [15–27]. Collectively, past research underscores the central role of ankle stiffness in shaping gait mechanics and functional walking performance.

Several studies have examined how ankle stiffness influences walking biomechanics using AFOs [6, 10, 28], with many focusing on the effect of stiffness on ankle *kinematics*. Generally, increasing AFO stiffness reduces peak ankle plantarflexion, peak dorsiflexion, and overall ankle range of motion [10]. Additionally, increased ankle stiffness has been associated with increased knee flexion at heel-strike, increased peak flexion, and decreased peak extension during stance [10].

In contrast, the effects of altered ankle stiffness on metabolic cost, muscle activation, and mechanics at proximal joints are still not well understood. Among studies investigating metabolic consequences [10], one notable example demonstrated that a purely passive mechanism could decrease the metabolic cost of level-ground walking at a constant speed [13]. This device provided unidirectional assistance, engaging a spring during stance and disengaging during swing. Building on this, the work of [29] applied unidirectional ankle stiffness in a similar manner and found a reduction in metabolic costs at multiple walking speeds, but not all, for certain stiffness values. In contrast, a study using a resistive device to modulate ankle stiffness bi-directionally found no significant metabolic benefit in children with cerebral palsy [30]. Studies examining the effects of stiffness on muscle activation have also been mixed [10, 13, 31–33], with one study reporting largely insignificant effects of altered ankle stiffness on lower-limb muscle activity [32]. The same study, however, found that knee joint work increased significantly with greater AFO stiffness [32]. Simulation results further suggest that an optimal bi-directional ankle stiffness may minimize hip joint work [28]. Despite these predictions, experimental studies have not reported significant effects of ankle stiffness on hip kinetics over the range of stiffness values explored in previous work [10]. To extend prior work, the present study more broadly examines how bi-directional stiffness applied in parallel with the ankle through a resistive device affects lower-limb kinetics, kinematics, and muscle activation in a healthy musculoskeletal system.

Conventional clinical AFOs, while highly useful in clinical applications, are not suitable for systematically investigating the effects of ankle stiffness on gait, as their stiffness cannot be easily adjusted in a controlled manner [28, 32, 34]. To overcome this limitation, we developed a passive ankle exoskeleton with interchangeable titanium springs that enabled us to systematically alter the stiffness and neutral angle across a targeted range of conditions. Compared to studies relying on commercial AFOs, in which different devices were required for different stiffness levels [17, 35], our single modular device provided consistent experimental conditions and allowed us to experimentally isolate the effects of ankle stiffness and neutral angle on gait dynamics. In addition to evaluating the impact of modifying ankle stiffness, we also varied the neutral angle because, under bidirectional stiffness, neutral angle and ground slope determine when during gait the spring stores and releases energy. This creates a potential trade-off between the energy required for deformation and the energy returned later in the gait cycle. We tested our device in healthy, young adults, to examine the effects of varying unilateral ankle stiffness within a healthy musculoskeletal system [31, 33, 36, 37]. Our central goal was to establish the fundamental relationship between ankle stiffness and gait mechanics in the absence of pathology, before extending these investigations to clinical populations. Using the ankle exoskeleton, we characterized the impact of varying equal bi-directional linear ankle stiffnesses on walking kinematics, kinetics, joint work, and muscle activation during level-ground, incline, and decline walking. These measurements allowed us to assess how individual joints and muscles of the affected leg compensate for changes in ankle power [38], as well as how stiffness alters the total mechanical energy and muscle activation of the limb. Understanding these effects may ultimately inform rehabilitation strategies by clarifying how altered ankle stiffness, whether caused by pathology or treatment, influences gait throughout the body.

## 2 Methods

### 2.1 Participants

This study was approved by the UCLA Institutional Review Board (protocol #23-001002). We recruited 9 able-bodied adults over 18 years old with no known pathologies that could affect gait (4 female, 5 male; mean body mass 72.35 *±* 12.05 kg; mean height 172 *±* 7 cm; mean age 25.6 *±* 2.7 years; mean Men’s US shoe size 8.8 *±* 1.5). After receiving a description of the experimental protocol, participants self-reported whether they had the ability to walk for 3 hours, wear an ankle exoskeleton, and complete the required locomotion tasks within a 5-hour time period. All subjects provided written informed consent prior to starting the experiment.

### 2.2 Passive Ankle Exoskeleton

We designed and built a passive ankle exoskeleton with interchangeable springs that allowed us to systematically vary ankle stiffness (Fig. 1(a)-(d)). The ankle exoskeleton frame was adapted from designs of [39, 40]. We manufactured interchangeable carbon fiber exoskeleton struts to fit 5 shoe sizes (Men’s US 7, 8, 9, 10, and 11), which we selected to accommodate most potential participants. Specifically, for each shoe size, we adjusted the lengths of the heel and toe struts such that the exoskeleton’s center of rotation aligned with the biological ankle. We also changed shank strut lengths for each participant, to ensure that the top straps secured comfortably below the knee for each user.

**Fig. 1.**
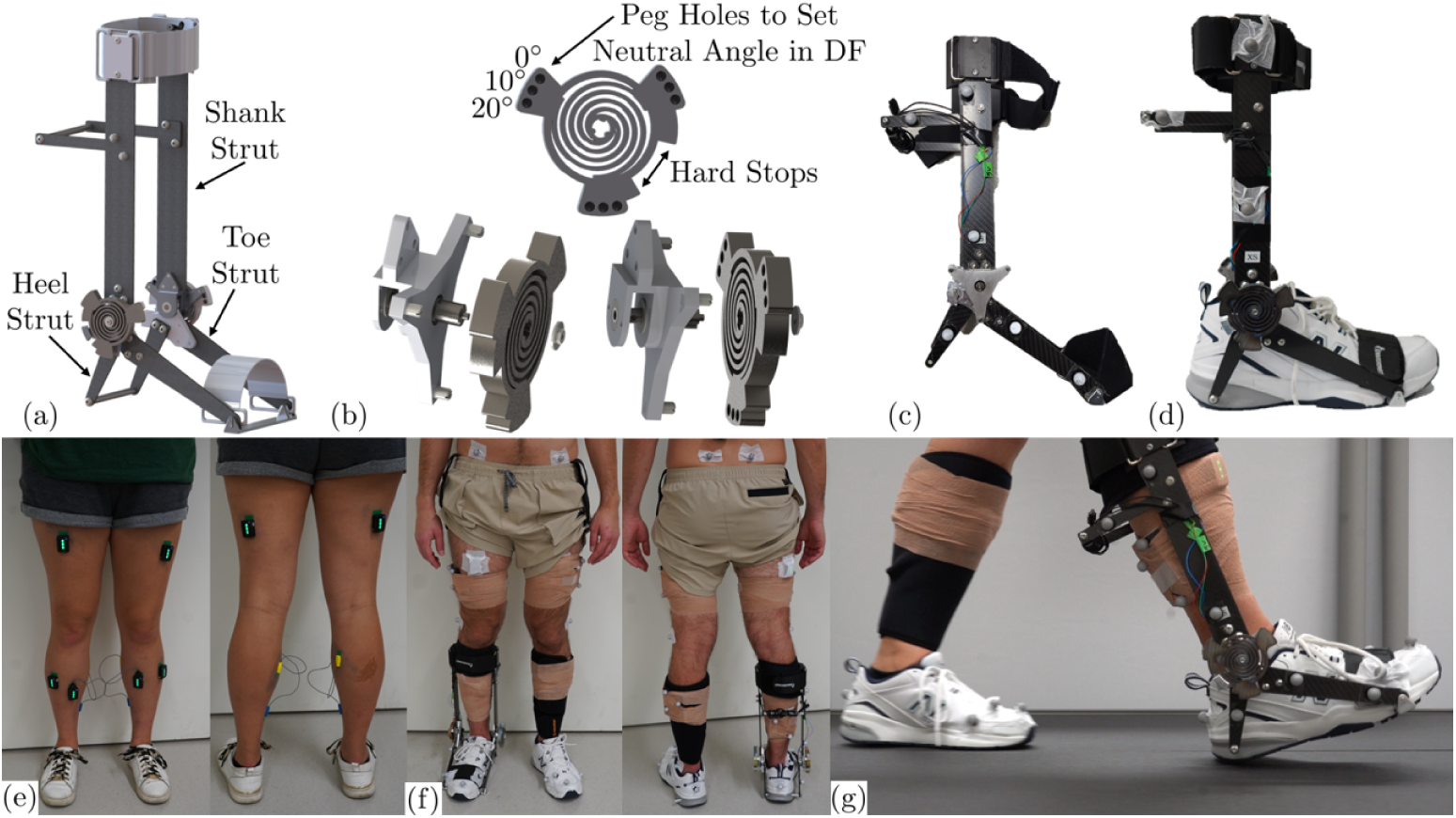
(a) Ankle exoskeleton CAD design with shank, heel, and toe struts. (b) Interchangeable spring and attachment mechanism with peg holes to change neutral angle in dorsiflexion direction and hardstops to enforce ROM. (c) Assembled exoskeleton with motion capture markers. (d) Exoskeleton attached to shoe. Heel struts are connected to a bar through the heel of the shoe and toe struts are connected to a toe plate placed in a cut-out of the shoe sole, secured with a strap across the toe of the shoe. (e) EMG sensors on a subject to record the rectus femoris, tibialis anterior, biceps femoris, gastrocnemius (medial), soleus. (f) Motion capture markers on the front and back of a subject. (g) Treadmill testing with exoskeleton and spring.

We incorporated torsional springs at the exoskeleton ankle joint to generate torque for motion past the neutral angle. We designed the device to enable quick spring changes by sliding them on and off pegs on the exoskeleton frame and securing with a nut. Additionally, each spring had multiple peg holes that could be used to set a range of neutral angles at 0 deg, 10 deg, or 20 deg in dorsiflexion. We used finite element analysis to design multiple planar coil springs by adjusting their coil thickness to achieve the desired target stiffnesses. In total, we manufactured 5 coil springs that yielded stiffness values of 10, 30, 50, 70, and 90 Nm/rad in simulation. This range of stiffnesses aligns with the range of stiffnesses of commonly used plastic AFOs [41]. Other studies have based their stiffness selection on these values and focused on applying stiffness in only one direction [42–44]. Our study differs in that we explore the effects of equal bi-directional stiffness. The springs were made from a medical-grade titanium alloy (Ti-6Al-4V). We experimentally characterized the actual stiffness of each spring by fixing the toe strut of the exoskeleton to a board, hanging weights from the calf strut, and measuring the angular displacement with an encoder (Table 1).

**Table 1.** Range of spring stiffnesses used for experiments. Target stiffness values were defined by finite element simulations, and actual stiffness values were measured experimentally. The ROM was restricted to *±*15.00° for stiffness values 10 - 70 Nm/rad. The ROM for the 90 Nm/rad spring was determined through simulations that showed this as the maximum allowable ROM for the prescribed stress limit.

| Target Stiffness | Actual Stiffness |  | ROM |
| --- | --- | --- | --- |
| Nm/rad | Nm/rad | Nm/deg | deg |
| 10 | 9.10 | 0.159 | $\pm 15.00$ |
| 30 | 27.57 | 0.481 | $\pm 15.00$ |
| 50 | 44.65 | 0.779 | $\pm 15.00$ |
| 70 | 74.17 | 1.294 | $\pm 15.00$ |
| 90 | 92.63 | 1.617 | $\pm 13.52$ |

We added hard-stops to the exoskeleton frame and on each spring to control the device’s range of motion (ROM) and prevent the springs from exceeding their maximum allowable stress. Simulations showed that the 90 Nm/rad spring could deflect by *±*13.52° without surpassing the prescribed stress limit, while the 70 Nm/rad spring could deform slightly past *±*15°. To standardize testing conditions with a range of springs, we constrained all softer springs to a maximum of a *±*15° ROM (Table 1). Although the 90 Nm/rad spring had a ROM limit of *±*13.52°, we opted to increase the ROM for other springs to *±*15° so as to not overly restrict the ankle when possible. Biological ankles move through about a 30° range during walking [45]), but prior studies have shown that restricting ROM to *±*15 deg does not significantly impact gait kinematics or joint work [46]. Therefore, while our study restricts both ROM and stiffness, we expect most changes in gait kinematics result from changes in stiffness rather than ROM.

### 2.3 Exoskeleton Fitting and Training

The exoskeleton was assembled for each participant based on their specified shoe size and with the respective carbon fiber struts. For all participants, New Balance 608 shoes were used (Fig. 1 (d)) and a calf wrap was worn on their non-exoskeleton leg to protect it from accidentally scraping the exoskeleton (Fig. 1 (g)). During each subject’s first visit, we assessed their self-selected walking speed (SSWS) by slowly increasing the treadmill speed until the subject indicated that the walking speed was too fast, then slowly decreasing the speed until the subject said the speed felt comfortable. This process was conducted for the level ground, incline (5°), and decline (−5°) walking conditions. The subjects were then fit with the exoskeleton and allowed an “adaptation period” during which they used the stiffest and softest spring at each walking condition until they reported feeling comfortable. Occasionally, the SSWS was adjusted during this adaptation period in response to the subject’s preference with the exoskeleton. The mean SSWSs for level ground, incline, and decline were 1.10 *±* 0.12, 0.99 *±* 0.12, and 1.01 *±* 0.14 m/s, respectively.

### 2.4 Experiments

During each participant’s second visit, they completed three-minute walking trials for all 31 walking conditions. Initially, two baseline conditions were tested. For the first baseline test, each subject walked on all three treadmill incline conditions (0°, 5°, −5°) without the exoskeleton. For the second, each participant donned the exoskeleton without a spring and walked at all inclines again, Fig. 1 (g). The exoskeleton was always worn unilaterally on the right leg. Following these 0 Nm/rad trials, spring configurations were varied randomly between walking trials. Specifically, the order of the spring walking trials was randomized by stiffness and neutral angle, while constraining incline trials to occur in groups and all trials of a given stiffness and neutral angle to be consecutive. Participants were blinded to spring conditions. All stiffness and neutral angle conditions were tested on level ground. A subset of stiffnesses and neutral angles were tested for the incline and decline conditions: stiffness values 10, 50, and 90 Nm/rad at 0° neutral angle and stiffness value 50 Nm/rad at 10° and 20° dorsiflexion neutral angle.

Kinematic data were collected with a 10-camera motion capture system (VICON, Oxford, UK) at 200 Hz using 14 mm markers, Fig. 1 (g) and (h). A 36 Cleveland Clinic marker model set was used (Supplementary Figure S1), modified to include 3 marker clusters on each of the exoskeleton shank and foot. Ground reaction forces were measured with a force-plate instrumented treadmill (FIT5, BERTEC, Colombus, OH) at 2 kHz. Electromyography (EMG) signals were measured from the biceps femoris, rectus femoris, tibialis anterior, soleus, and gastrocnemius (medial) on both legs using wireless EMG sensors (Trigno, Delsys, Natick, MA) at 2 kHz (Fig. 1 (e) and (f)). Before EMG sensor placement, all subjects removed body hair from the areas of skin where EMG sensors were to be placed. EMG sensors were placed according to SENIAM guidelines [47].

### 2.5 Data Analysis

To minimize the effects of adaptation to each new condition in the analysis, only the last minute of each three-minute walking trial was analyzed. Motion capture marker data were labeled and gap filled in Vicon Nexus software. These data were then filtered by a 15 Hz low-pass Butterworth filter. Ground reaction force data were filtered with a 3 Hz low-pass Butterworth filter and downsampled to 200 Hz. These data were also rotated for the incline and decline trials to reflect the rotation of the force plates. This degree of rotation was analytically determined from the marker clusters on the treadmill in the incline/decline position relative to the level-ground position.

A subject-specific model was created using the OpenSim *gait10dof18musc* base model and the static standing trial collected for each participant. A second model was created for each subject for the trials that used the exoskeleton. The exoskeleton mass and inertial properties were determined from a CAD model of the exoskeleton and added to the tibia and calcaneus bodies of the subject model on the leg that wore the exoskeleton. These subject-specific scaled models, with all joints unlocked, were used in OpenSim to compute inverse kinematics and inverse dynamics from the motion capture data and ground reaction force data. The joint angles obtained through inverse kinematics were baseline normalized by subtracting the mean joint angle from the respective static trial, such that a joint angle of 0 deg represents quiet standing. Exoskeleton joint angle was computed analytically (separate from the OpenSim inverse kinematics) based on single-value decomposition of the marker clusters fixed to the exoskeleton frame. The exoskeleton angles, experimentally characterized spring stiffness, and spring neutral angle were used to calculate the moment and power of the exoskeleton joint. To determine the net biological ankle moment of the subject’s exoskeleton-side ankle joint, the exoskeleton moment was subtracted from the net total ankle moment obtained through inverse dynamics.

All EMG data were filtered using a bandpass filter with a passband between 20 and 500 Hz, fullwave rectified, and integrated using a 50 ms moving average filter. Integrated EMG data from each electrode were normalized to the maximum EMG value recorded by that electrode from the level-ground walking trial without the exoskeleton. Sensor errors were found in the data for 3 subjects. For one subject, the EMG data was erroneous across all trials, showing muscle activation patterns with no resemblance to typical human gait; this subject’s EMG data were excluded from analysis. For two other subjects, this error only existed in the 0 Nm/rad level ground stiffness trial; EMG data from these trials was excluded from analysis. Additionally, an error was found in our experimental setup for one subject for the 30 Nm/rad stiffness trials for all neutral angles and the 10 Nm/rad stiffness incline trial at 0° neutral angle. Since these errors were found after the subject’s second experimental day, this subject came in for a third experimental day to redo these tests. The baseline trials without the exoskeleton were repeated and the maximum EMG values from the level-ground trial were used to normalize the EMG data from the trials completed that day.

Kinematic, kinetic, and EMG data for each trial were segmented by stride, based on the timing of consecutive heel-strikes on the exoskeleton side. The data were also segmented by exoskeleton-side stance phase and swing phase, determined by the timing of exoskeleton-side heel-strike to toe-off and vice versa. The heel-strike and toe-off timing were determined by thresholds on the vertical ground reaction force of the exoskeleton side. Each data segment was normalized to 0-100% gait phase.

To assess the effects of ankle stiffness on the energy output of each subject, we considered two different metrics: mechanical energy and muscle activation. To quantify the mechanical energy, we first calculated the instantaneous power at each joint in the sagittal plane as the product of the instantaneous joint velocity (numerically differentiated joint angle and time) and the instantaneous joint moment. We then numerically integrated the joint power during the times of positive power, negative power, and over the entire gait phase to obtain the positive, negative, and net work of the joint, respectively[48]. Summing the work of the hip, knee, and ankle of the exoskeleton-side gave measures of the total positive, negative, and net mechanical energy of the affected leg. The mean muscle activation across each gait phase was calculated for every muscle for each subject. This mean activation was averaged across the 5 exoskeleton-side muscles to provide a measure of the mean muscle activation per affected-side muscle. Kinematic asymmetry of each joint was quantified as the root mean square difference (RMSD) between the left and right leg mean sagittal-plane joint kinematics. We obtained mean trajectories by averaging time-normalized strides (heel-strike to heel-strike of the respective leg) within a trial for each subject. The three joint RMSDs were averaged to measure the mean kinematic asymmetry per joint, as an indicator of impact on full-leg kinematic symmetry.

### 2.6 Statistical Analysis

All statistical analyses were conducted in MATLAB ([R2022a], MathWorks, Natick, MA). The goal of our statistical analysis was to assess the effect of stiffness on the following measures: kinematic asymmetry, mechanical energy (positive, negative, and net), mean muscle activation, and peak muscle activation. We conducted repeated measures of analysis of variance (ANOVA) tests to assess the effect of ankle stiffness on the aforementioned measures. Where the ANOVA showed a statistically significant (p *<* 0.05) effect of stiffness, we conducted a post-hoc Tukey’s Honestly Significant Difference (HSD) to make direct pairwise comparisons between stiffnesses (*α <* 0.05). We also created first- and second-order linear mixed effects (LME) models using MATLAB *fitlme* to evaluate whether a significant linear or quadratic relationship existed between spring stiffness and each measure of interest. In these models, we modeled spring stiffness as the fixed effect and included the subject as a random intercept. For the first-order models, the linear fixed-effect parameter of stiffness was evaluated for significance. Linear and quadratic stiffness terms were included in the second-order models and only the quadratic parameters were evaluated for significance. For both models, we set significance levels to 0.05.

We performed these statistical tests for two ranges of stiffness: 0-90 Nm/rad and 0-70 Nm/rad. The springs in this second range had a uniform ROM, compared with the first range where the 90 Nm/rad spring had a smaller ROM. The experimentally-derived stiffness values (Table 1), rather than the idealized target stiffnesses, were used for all statistical analyses.

### 2.7 Sample Size Justification

We performed a power analysis to justify the sample size for detecting subject-specific changes in joint mechanical work across stiffness conditions. We performed this power analysis based on pairwise comparisons of joint work between stiffness conditions; this metric was chosen because it is the study’s primary mechanical outcome and directly reflects redistribution of mechanical work throughout the leg in response to altered ankle stiffness. Based on an *a priori* power analysis framework for a two-sided paired comparison (*α* = 0.05), our sample size of *n* = 9 participants provided 80% power to detect a paired difference of 0.630 J/kg, equivalent to a 3.697% change in work relative to the reference condition and a standardized effect size Cohen’s *dz* = 1.067. This sensitivity analysis supports the adequacy of the sample size for detecting meaningful subject-specific effects in joint work across stiffness conditions.

## 3 Results

### 3.1 Joint Angles, Net Joint Moments, and Joint Power in Level Walking

All subjects were able to walk while wearing the exoskeleton at all tested stiffnesses (Fig 2). Wearing the exoskeleton with no spring had little impact on ankle angle, moment, or power. Ankle range of motion decreased progressively with increasing spring stiffness (Fig 2, left). This effect was primarily attributable to a decrease in the maximum plantar flexion ankle angle immediately after toe-off, with a small progressive reduction also observed in the peak stance-phase dorsiflexion angle. Although spring deflection generally decreased with increasing stiffness, the magnitude of the spring’s resistance torque (i.e. deflection *×* stiffness) increased with increasing stiffness. There was a concomitant reduction in ankle joint moment during powered plantar flexion as the spring torque increased with increased spring stiffness. Peak power (both positive and negative) at the biological ankle joint decreased with increasing exoskeleton stiffness (Fig 2, right).

**Fig. 2.**
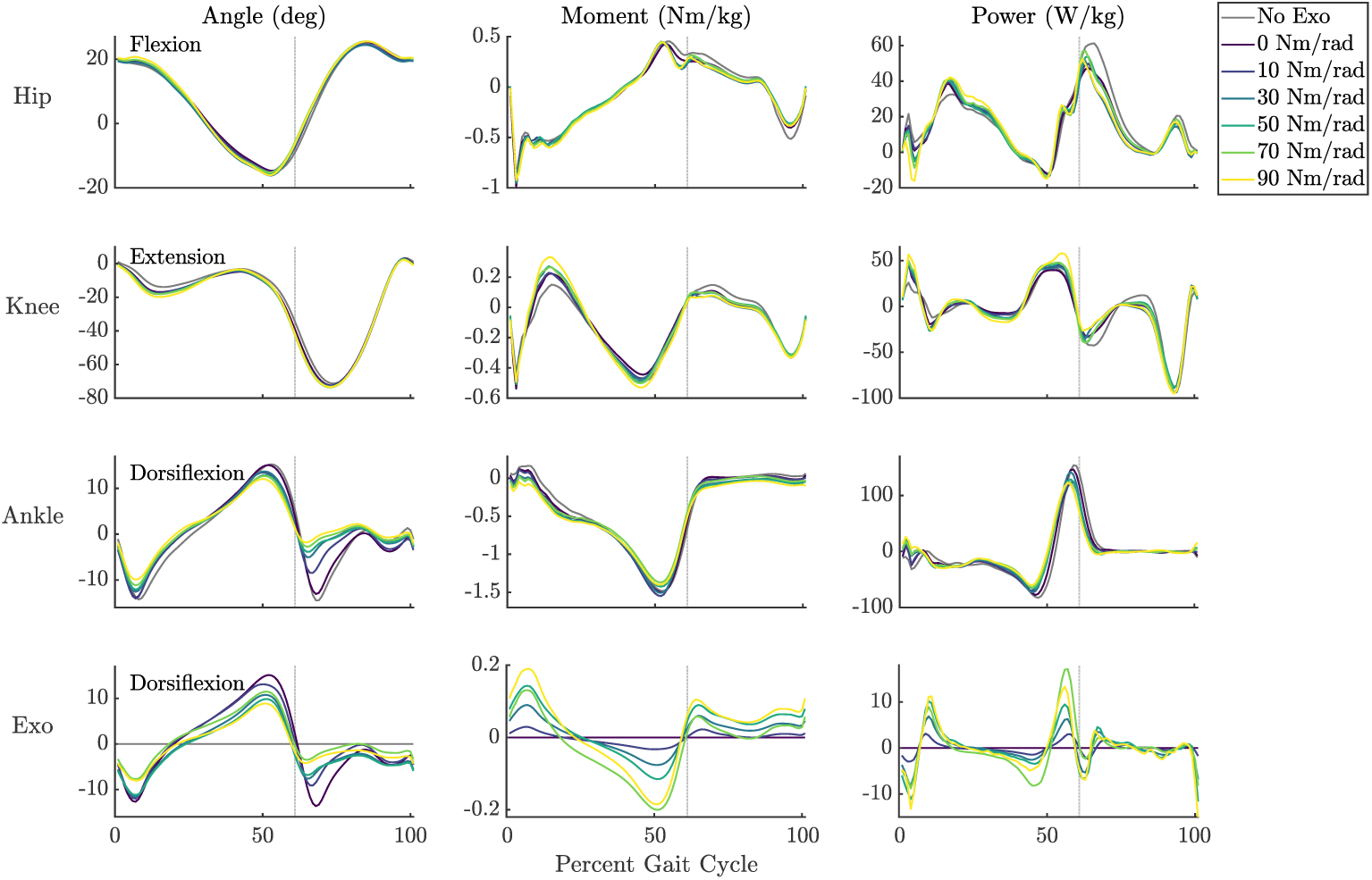
Kinematics (left), kinetics (middle), and power (right) of exo-side joints for different spring stiffness conditions on level ground walking with 0° neutral spring angle. Gray dashed line represents the approximate stance to swing transition. Biological kinematics (hip, knee, and ankle) were computed from motion capture data using inverse kinematics, while the exoskeleton kinematics were determined with singular-value decomposition of exoskeleton marker clusters. Exoskeleton moments and power were calculated based on these kinematics and the spring stiffness. Net joint moments were obtained from motion capture data and ground reaction force data using inverse dynamics. The biological ankle moment was determined by subtracting the exoskeleton moment from the net ankle moment. Lines represent intersubject mean trajectories for different spring conditions. Joint angles are reported relative to quiet standing.

The addition of stiffness in parallel with the ankle also affected the dynamics of the knee and hip, most noticeably in joint moments and powers (Fig 2, middle and right). Peak knee moment increased with increasing spring stiffness during both early stance flexion and mid stance extension. Magnitude of knee power tended to increase across the gait cycle as the spring stiffness increased, except just after toe-off when the stiffest spring resulted in the least-negative knee joint power. Trends in hip moments and powers were less pronounced, with trade-offs between spring stiffnesses across the gait cycle.

### 3.2 Kinematic Symmetry

Stiffer springs resulted in more kinematic asymmetry in all three joints (Fig 3). The relationship between mean kinematic RMSD across all three joints and spring stiffness was significantly linear (1st order LME, p *<* 0.001) in the range of 0 to 90 Nm/rad. This trend was seen in each individual joint for this range as well (LME p *<* 0.016). In the range of 0 to 70 Nm/rad, quadratic trends were present for the knee, ankle, and mean RMSD across the three joints (2nd order LME p *≤* 0.009). The maximum kinematic asymmetry of the data in this range occurred at 30 Nm/rad, while the maximum of the LME occurred around 50 Nm/rad. All LME and ANOVA results for kinematic RMSD are provided in Supplementary Tables S1 and S2, respectively.

**Fig. 3.**
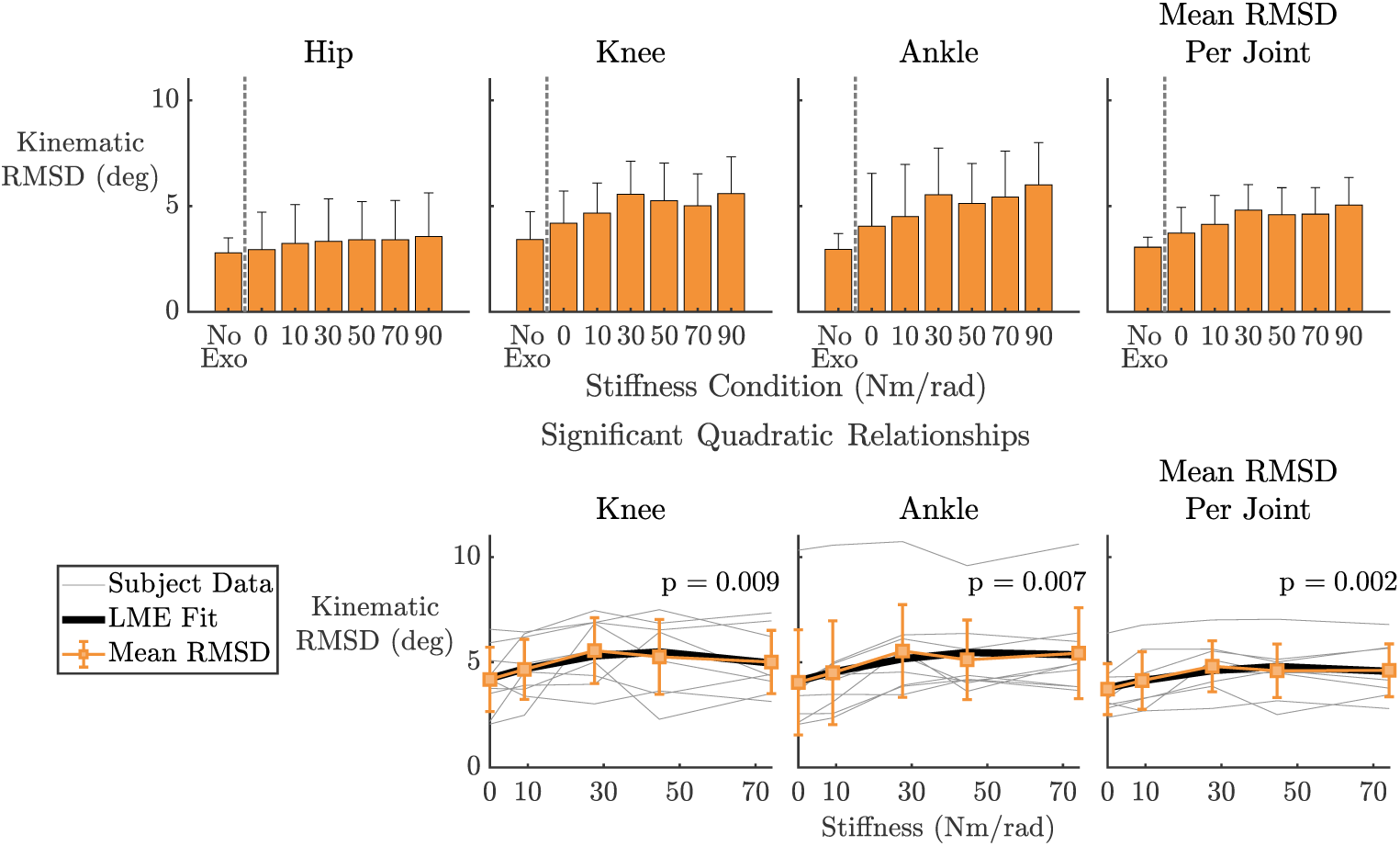
(Top) Kinematic RMSD between left and right side joints, showing the degree of kinematic asymmetry with respect to spring stiffness. Mean RMSD per joint is calculated as the mean of the individual joint RMSDs. The bars indicate the intersubject mean and the error bars represent intersubject standard deviation. (Bottom) Linear mixed effect model results (black) across the control condition (0 Nm/rad) and spring conditions with an equal ROM (*±*15 deg). These models show the significant quadratic relationships (p *<* 0.05) between kinematic RMSD and spring stiffness with their respective p-values. The intersubject mean RMSD with standard deviation (orange) and mean RMSD data for each subject (gray) are also given.

### 3.3 Joint Mechanical Work

Total net work across all three joints increased linearly (1st order LME, p *<* 0.001) with spring stiffness (Fig. 4 bottom right). This effect was dominated by a linear decrease in net work at the ankle joint (p *<* 0.001), predominantly during the stance phase. Negative work at the knee joint across the full gait cycle increased linearly with stiffness (p = 0.047). This was driven primarily by changes during the stance phase, during which negative knee work was lowest in the zero stiffness condition, and significantly higher for all stiffnesses above 10 Nm/radian (ANOVA with posthoc Tukey HSD, p *<* 0.035 for all relationships). However, this increase in negative work was offset by a linear increase in positive knee work (1st order LME, p = 0.004), such that there was not a significant relationship between stiffness and network at the knee joint (p = 0.085). Network at the hip joint was significantly affected by stiffness (ANOVA, p = 0.027). In the range of stiffnesses from 0 to 70 Nm/rad, net work at the hip was significantly quadratic (2nd order LME, p = 0.005), with a minimum near 30 Nm/rad. Examining hip power during the gait cycle links this phenomenon to early swing phase (Fig. 2). Network at the hip joint was significantly lower at 30 Nm/rad than at zero stiffness (Tukey HSD, p = 0.005). All LME and ANOVA results for joint mechanical work are provided in Supplementary Tables S1 and S2, respectively.

**Fig. 4.**
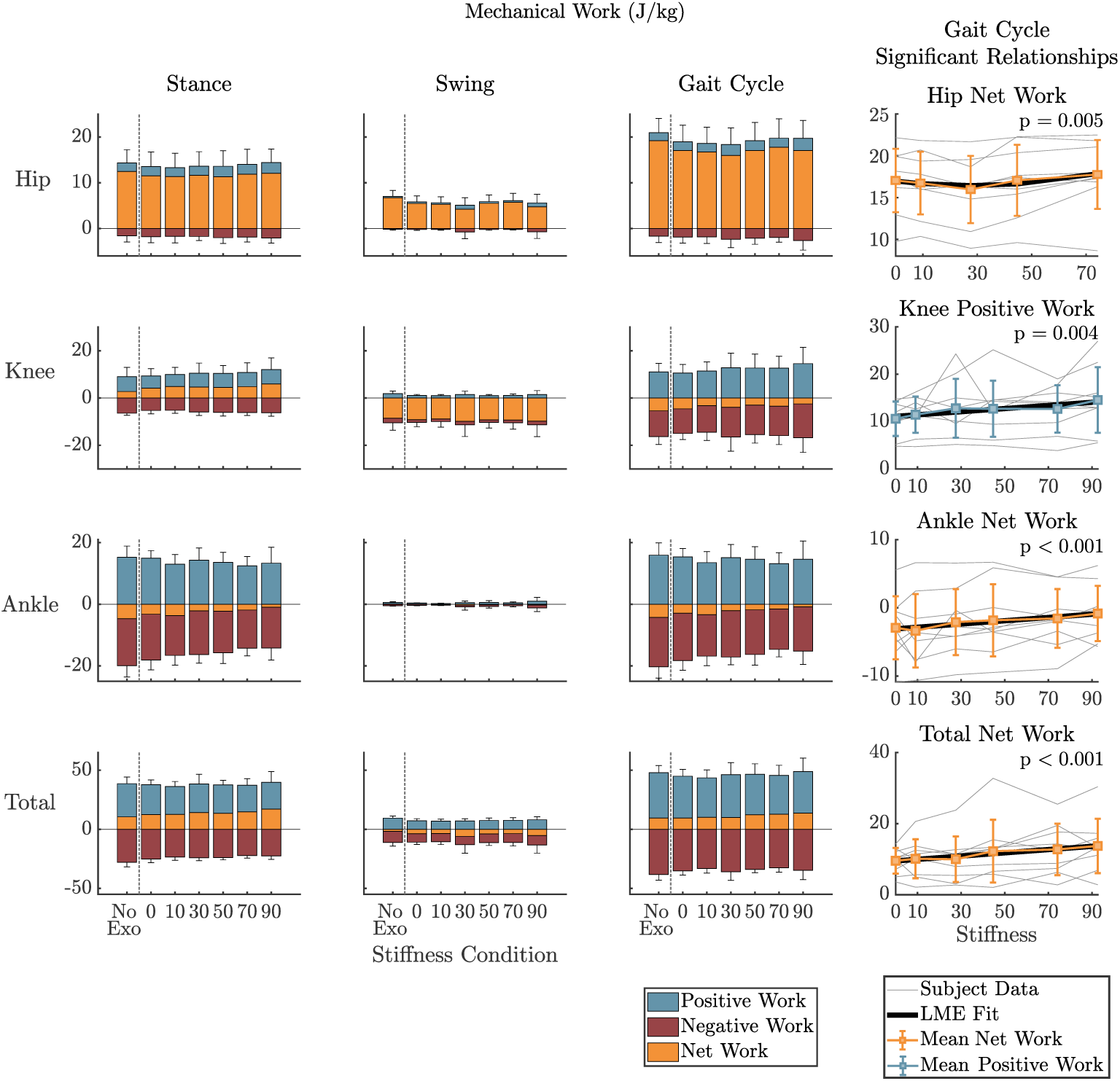
Mean mechanical work of exo-side joints for stance (left), swing (middle-left), and the full gait cycle (middle-right). (Bottom row) Total mechanical energy of exo-side joints. Bar represent intersubject means, and error bars show intersubject standard deviation. (Right) Linear mixed effect model results (black) for the gait cycle with their respective p-values. A significant quadratic relationship (p *<* 0.05) for the hip network was found over the control condition (0 Nm/rad) and springs with an equal ROM (*±*15 deg). Significant linear relationships (p *<* 0.05) were found for the knee positive work, ankle network, and total net work with respect to stiffness across the full range of spring conditions. The intersubject mean network and positive work with standard deviations are shown in orange and blue, respectively. Individual subject means are shown in gray.

### 3.4 Muscle Activation

Peak activation of the tibilias anterior (1st order LME, p *<* 0.001) and soleus (p *<* 0.001) showed a significant inverse linear relationship with spring stiffness, as did mean soleus activation (p *<* 0.001). Mean activation across all leg muscles showed a significant quadratic relationship with stiffness from 0 to 70 Nm/rad (2nd order LME, p = 0.021), with the lowest measured mean activation at 30 Nm/rad. The mean rectus femoris activation exhibited a quadratic trend with spring stiffness, with a minimum at 30 Nm/rad, however this trend was not significant (p = 0.056). All LME and ANOVA results for muscle activation (mean and peak) are provided in Supplementary Tables S1 and S2, respectively.

### 3.5 Slope Walking and Spring Neutral angle

As with level ground walking, spring stiffness has a substantial effect on ankle joint kinematics during incline and decline walking (Fig. 6). Across terrain conditions, the absolute effect of stiffness on ankle motion for a given portion of the gait cycle (e.g. early stance phase) appears to scale with the range of ankle motion during that part of the gait cycle. For example, the absolute effects of spring stiffness during controlled plantar flexion in early stance (immediately following heel-strike) are smaller for incline walking than for decline or level-ground walking (Fig. 6, top row), likely due to the diminished range of the ankle during those terrain conditions. Similarly, the effects of spring stiffness during early swing (immediately following toe-off) are smaller for decline walking than for incline or level-ground walking (Fig. 6, top row).

The effects of spring neutral angle were most pronounced during controlled plantar flexion in early stance, and in the kinematics throughout the swing phase (Fig. 6, middle column). More-dorsiflexed neutral angles led to less plantar flexion during these parts of the gait cycle, with greater effect as stiffness increased. For all three terrain conditions, ankle kinematics during early stance and throughout swing were closest to the “no exo” case at an neutral angle of 0 degrees (Fig. 6). Peak dorsiflexion angle during stance increased with increasing neutral angle across stiffnesses and terrain conditions, converging on the kinematics of the “no exo” case. However, the magnitude of this effect was small compared to the effect of both terrain and neutral angle during early stance and throughout swing (Fig 6).

Because of the hard-stops on the spring, changing the neutral angle inherently also shifted the ROM, meaning the results identified here are a function of both the spring stiffness and ROM. However, the absence of noticeable flat-line periods in the ankle kinematics near its ROM limits indicates that there was not prolonged contact with the hard-stops in the experiments. This suggests that the limited ROM did not significantly alter the ankle kinematics and the main effects observed are primarily related to the spring stiffness.

## 4 Discussion

In this work, we studied the impact of linear bi-directional stiffness applied in parallel with the ankle on lower-limb gait biomechanics using a passive ankle exoskeleton we designed with interchangeable springs. We analyzed level ground, incline, and decline gait biomechanics using traditional motion capture, force plates, and surface EMG. Given ankle stiffness can be altered by mechanical injury or assistive devices, the results of this work could inform rehabilitation by providing insight into how injury and treatment methods could affect an individual’s gait. This study showed that altered ankle stiffness can lead to changes in muscle activation and a redistribution of joint work. Recognizing the potential for overloading or underutilizing certain muscles and joints could inform intervention and treatment strategies to prevent further injury or deconditioning.

The results of this study suggest that conditions that increase bi-directional ankle stiffness can be expected to decrease ankle ROM and typically increase asymmetry of all joints of the ipsilateral limb, due primarily to decreased plantarflexion at toe-off. These results are consistent with past findings [10]. Ankle muscle activation can also be expected to decrease with increased ankle stiffness (Fig. 5). In this study, increasing ankle stiffness corresponded to linearly decreasing net ankle work over the gait cycle. As the stiffness increased, the dynamics of the spring increasingly dominated the ankle’s dynamics, causing the ankle’s network to tend towards zero. This aligns with the expected network of a spring over a cycle.

**Fig. 5.**
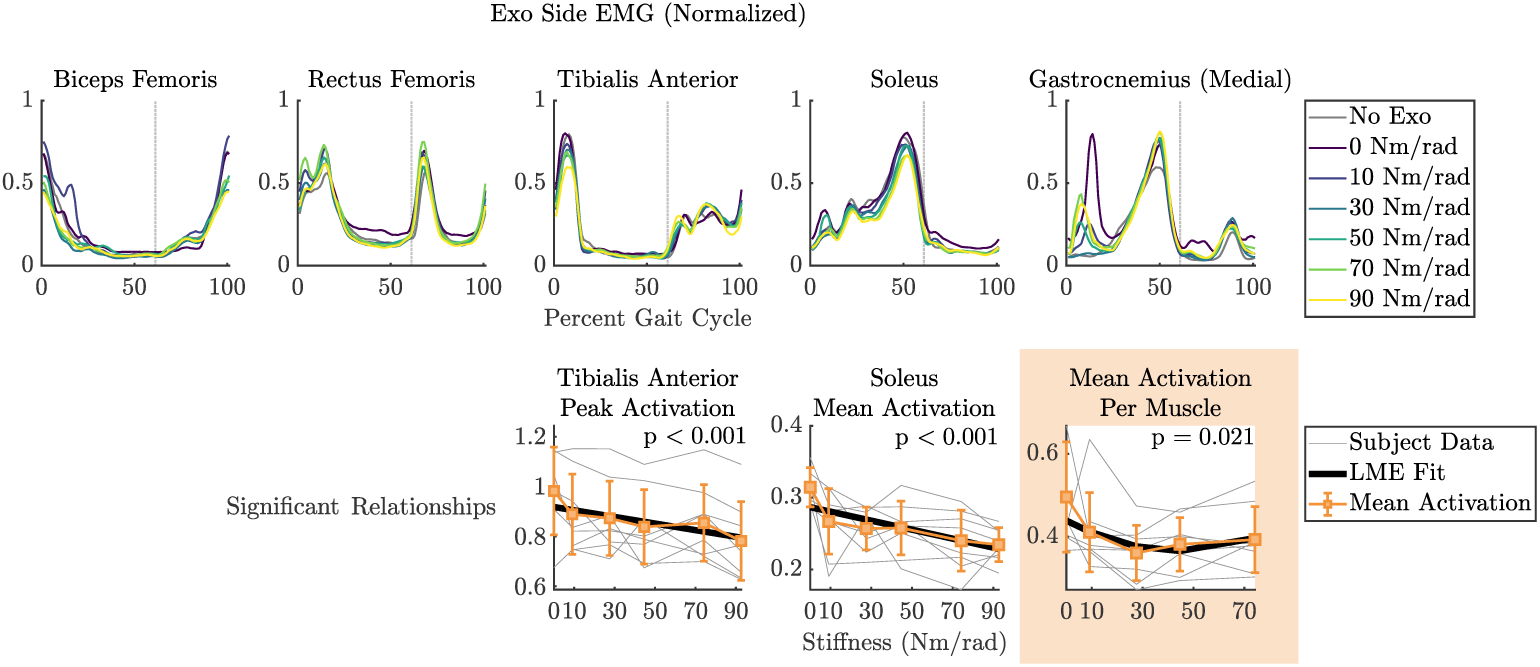
(Top) EMG activation for exo-side muscles for all spring stiffness conditions. Lines represent intersubject mean activation. Gray dashed line represents the approximate stance to swing transition. (Bottom) Linear mixed effect model results (black) with their respective p-values. Significant linear relationships (p *<* 0.05) were found for the tibialis anterior peak activation and soleus mean activation over the full range of spring stiffesses. A significant quadratic relationship (p *<* 0.05) for the mean activation per muscle was found across the control condition (0 Nm/rad) and springs with an equal ROM (*±*15 deg). The intersubject mean and standard deviations of each of these quantities are shown in orange, and the mean data for each subject are shown in gray.

**Fig. 6.**
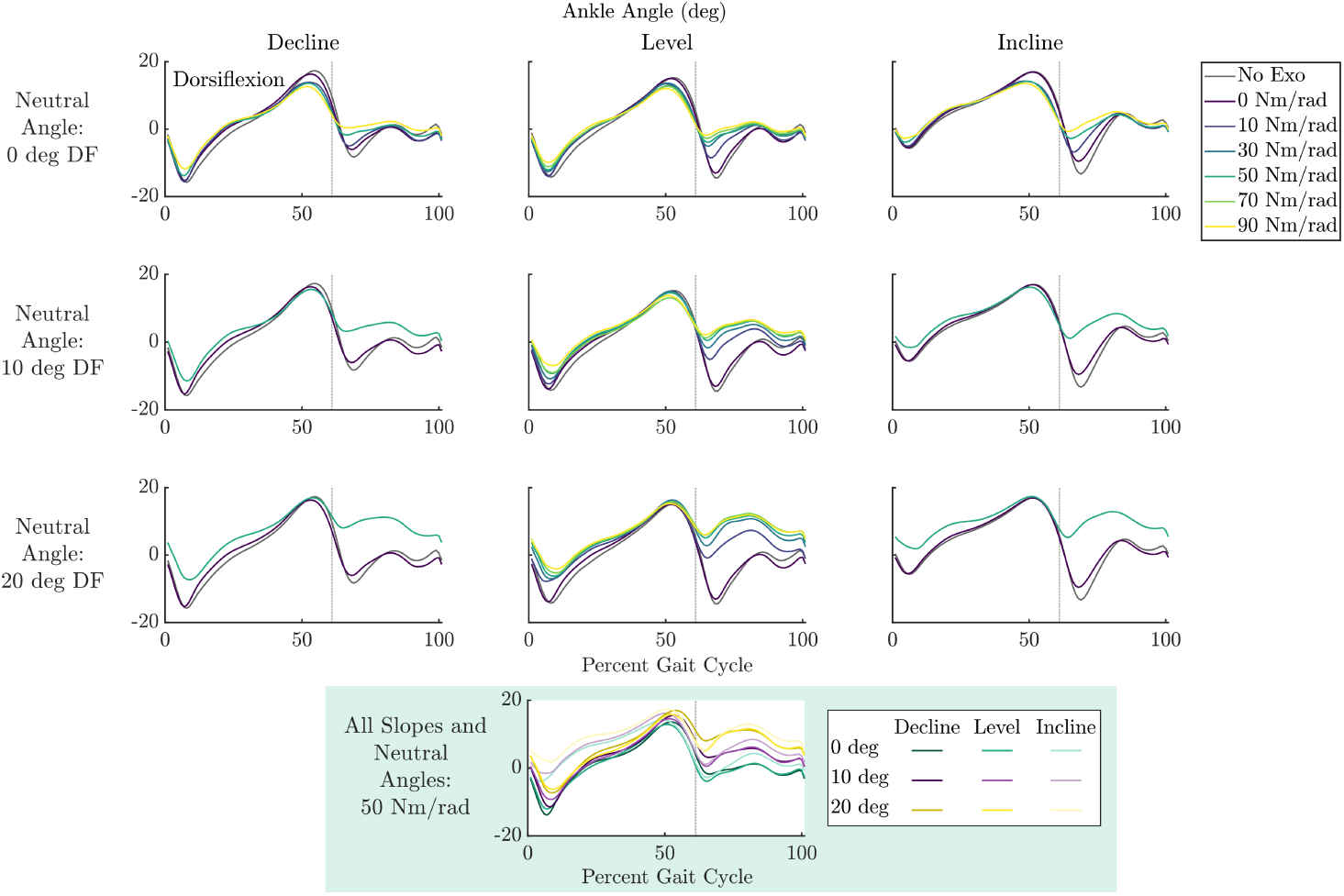
Ankle kinematics of all stiffness, dorsiflexion (DF) neutral angle, and incline conditions. Kinematics were obtained from motion capture data via inverse kinematics. (Bottom) Ankle kinematics for all incline and neutral angles conditions with the 50 Nm/rad spring. Lines represent intersubject mean trajectories. Gray dashed line represents the approximate stance to swing transition.

While some gait measures vary monotonically with ankle stiffness, others exhibit more complex, non-monotonic behavior, as reflected by local maxima and minima observed in this study. Specifically, over the 0-70 Nm/rad stiffness range, we observed maximum values for kinematic asymmetry of the knee, ankle, and values per joint with the 30 Nm/rad spring. Over this stiffness range, this spring also led to minimum values for hip joint net work and mean muscle activation on the exoskeleton-side leg. These minimum values at the 30 Nm/rad stiffness condition were even lower than the 0 stiffness condition. The reduction in hip joint network was present across most individual participants’ data (Fig. 4, top right). These findings imply that participants may have adopted an asymmetric adaptation to leverage the 30 Nm/rad spring in reducing mechanical energy expenditure at the exoskeleton-side hip joint. Additionally, the results reveal a tradeoff: decreasing hip energy and average muscle activation come at the expense of increasing kinematic asymmetry. Interestingly, the stiffness value that led to minimum work and and muscle activation when applied bidirectionally in this study is close to the stiffness value of 50 Nm/rad that led to minimum metabolic rate when applied unidirectionally in [29], where 50 Nm/rad was the lowest stiffness value tested. More work will be needed to understand why 30 Nm/rad specifically seems to be energetically favorable when applied bidirectionally in this population. However, this provides evidence that there may exist specific stiffness ranges that minimize joint work.

Many of the significant quadratic relationships observed in this study, such as those in kinematic asymmetry, net hip joint work, and mean muscle activation as a function of ankle stiffness, were only significantly quadratic in the range from 0 to 70 Nm/rad spring stiffness. Although this may be tied to the uniquely smaller range of motion for the 90 Nm/rad spring, it is also possible that the observed trends are part of a local effect over a limited stiffness range. One previous study found no significant impact stiffness on hip joint work when comparing between individuals’ prescribed AFO device and devices manufactured to be 20% more and less stiff [32]. This suggests that optimality may exist in a region of lower stiffness, where the ankle is also allowed greater range of motion during gait, and that these effects are likely masked by the dominant linear trends that emerge in stiffer devices. Additionally, among the previous studies that examined the impact of ankle stiffness on muscle activation, few significant effects were found between ankle stiffness and muscle activation [10, 32]. This provides further evidence that the low-stiffness regime is of particular interest when seeking minimum joint work and muscle activation.

The significant relationship between hip work and ankle stiffness found in this study is a promising result for lower-limb prosthesis research [49]. This phenomenon could be exploited to address the increased hip power expenditure in lower-limb prosthesis gait compared to that of able-bodied individuals [50]. By optimizing work of active prosthetic ankles, hip work could be reduced to avoid overtaxing the hip joint and impeding energy efficiency in walking [51].

The objective of this study was to understand the impact of unilateral bi-directional ankle stiffness on gait in able-bodied individuals. In this population, we found interesting phenomena centered at stiffnesses in the lower range of those typically prescribed to individuals who use AFOs. Our results shed new light on the ways in which people may adapt their gait to synergize with a low-stiffness spring in parallel with the ankle joint, which has the potential to inform the design of wearable devices for people without gait pathology. To fully understand how such devices may affect gait across terrains, further analyses could be performed on the impact of non-neutral spring set point and the effect of spring stiffness on inclines/declines along with transverse and frontal plane motion. Additionally, because our experiments were focused on steady-state walking, it is possible that certain stiffness conditions would be more or less favorable in events such as gait acceleration/deceleration, stair climbing, or obstacle avoidance. We also expect that this work will motivate future extension of these studies to individuals with pathologies affecting the ankle (e.g. stroke, paralysis), with specific emphasis on understanding how ankle stiffness impacts mechanical energy and muscle activation across all joints of the affected leg in these populations. Results showing similar effects to those identified in this study would be of particular interest to clinicians looking to prescribe treatments such as AFOs or other assistive devices.

## Acknowledgments

We thank Sara Meschi, Kaushal Patel, Leonardo Ruffini, and He Kai Lim for assisting with the exoskeleton design and manufacturing process. We appreciate Rasheedat Ekiyoyo’s assistance with the experiments. We also thank Deema Totah for her discussions on the analysis and the significance of the study in the context of the field of AFOs.

## Declarations

### CRediT authorship contribution statement

RGH: Conceptualization, Methodology, Formal Analysis, Investigation, Data Curation, Writing - Original Draft, Writing - Review & Editing, Visualization, Project administration. MJR: Methodology, Investigation, Writing - Original Draft, Writing - Review & Editing. WF: Methodology, Investigation. LH: Data Curation. AT: Methodology. ASV: Conceptualization, Writing - Review & Editing. TRC: Conceptualization, Writing - Original Draft, Writing - Review & Editing, Supervision.

### Supplementary Information

The online version contains supplementary material.

### Data Availability

The data that support the findings of this study are not openly available due to the data sharing restriction in the UCLA IRB protocol #23-001002 and the approved informed consent form. Data are located in controlled access data storage at the University of California, Los Angeles, and can be shared from the authors upon reasonable request.

### Ethical Approval

This study was approved by the UCLA Institutional Review Board and the methods were conducted according to the IRB-approved protocol #23-001002. All subjects provided their informed consent under this protocol before any subject data were collected.

### Declaration of generative AI and AI-assisted technologies in the manuscript preparation process

During the preparation of this work the authors used ChatGPT (OpenAI) to assist with editing and refinement of individual statements. The authors reviewed and revised all AI-assisted content and take full responsibility for the content of the published article.

