## Supplementary Material for "Effects of Ankle Stiffness on Total Leg Kinematics, Mechanics, and Muscle Activation during Walking"

### Supplementary Information

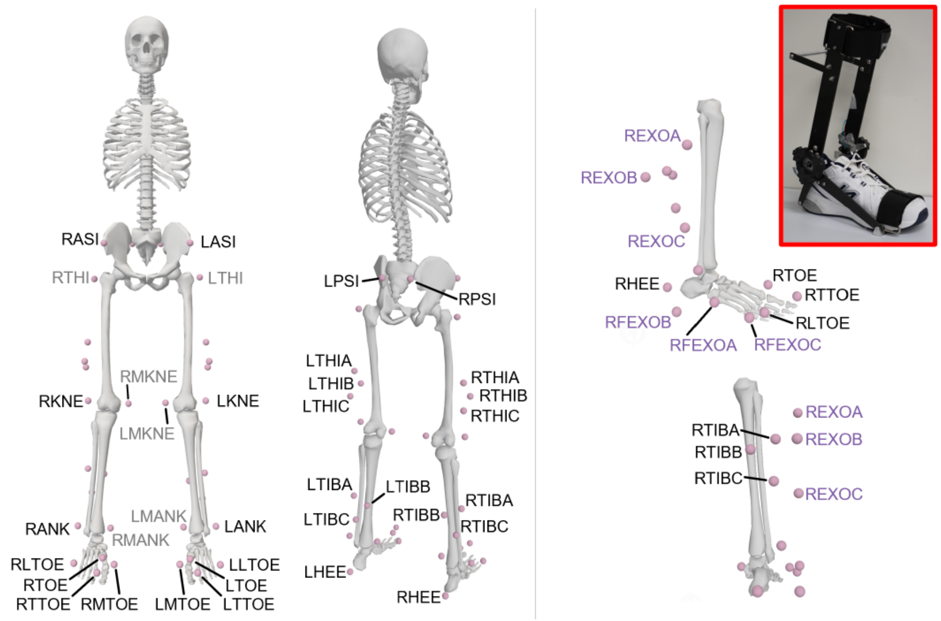


(a) (b)

**Fig. S1** OpenSim skeletal marker model set used for data collection. (a) A modified 36 Cleveland Clinic marker model set. Marker names in gray were used for the static scaling trial only and removed before walking experiments. (b) Six markers were added to this set to create a 3 marker cluster on both the exoskeleton shank and foot, as shown in the top right.

**Table S1** LME P-values for Gait Cycle

| Name | Linear 0-70 | Linear 0-90 | Quad 0-70 | Quad 0-90 |
|---|---|---|---|---|
| Hip Flexion RMSE Mean | **0.0413** | **0.0079** | 0.1786 | 0.3419 |
| Knee Angle RMSE Mean | 0.0778 | **0.0161** | **0.0091** | 0.1344 |
| Ankle Angle RMSE Mean | **0.0001** | **<0.0001** | **0.0066** | 0.0973 |
| Avg Joint RMSE Mean | **0.0015** | **<0.0001** | **0.0021** | 0.0709 |
| Hip Flexion Work Net Mean | **0.0367** | 0.1097 | **0.0047** | 0.4816 |
| Knee Angle Work Net Mean | 0.3619 | 0.0855 | 0.4159 | 0.8537 |
| Ankle Angle Work Net Mean | **0.0121** | **0.0004** | 0.5309 | 0.7580 |
| Total Work Net Mean | **0.0049** | **0.0001** | 0.9795 | 0.9804 |
| Hip Flexion Work Pos Mean | **0.0089** | **0.0013** | **0.0477** | 0.2993 |
| Knee Angle Work Pos Mean | 0.0726 | **0.0042** | 0.1679 | 0.7659 |
| Ankle Angle Work Pos Mean | 0.1234 | 0.4921 | 0.3545 | 0.7657 |
| Total Work Pos Mean | 0.3211 | **0.0352** | 0.3367 | 0.8253 |
| Hip Flexion Work Neg Mean | 0.5334 | **0.0198** | **0.0171** | 0.7711 |
| Knee Angle Work Neg Mean | 0.1775 | **0.0470** | 0.3741 | 0.8300 |
| Ankle Angle Work Neg Mean | **0.0003** | **0.0024** | 0.7189 | 0.5955 |
| Total Work Neg Mean | 0.1952 | 0.5713 | 0.3266 | 0.7977 |
| Bicep Fem Total Mean | 0.1579 | 0.0700 | 0.3083 | 0.2890 |
| Rec Fem Total Mean | 0.9043 | 0.4798 | 0.0563 | 0.5891 |
| Ta Total Mean | 0.4750 | 0.0634 | 0.1546 | 0.8128 |
| Soleus Total Mean | **0.0002** | **<0.0001** | 0.0677 | **0.0435** |
| Gas Med Total Mean | 0.9199 | 0.9210 | 0.1511 | 0.2605 |
| Bicep Fem Peak Mean | 0.1913 | 0.0912 | 0.5257 | 0.4790 |
| Rec Fem Peak Mean | 0.3196 | 0.8886 | 0.3139 | 0.7320 |
| Ta Peak Mean | **0.0145** | **<0.0001** | 0.0514 | 0.5759 |
| Soleus Peak Mean | **0.0130** | **0.0002** | 0.2225 | 0.4546 |
| Gas Med Peak Mean | 0.8453 | 0.9764 | 0.0876 | 0.1895 |
| Mean Activation Per Muscle Mean | 0.1696 | **0.0372** | **0.0210** | 0.1081 |

LME p-values for the gait cycle for both linear and quadratic relationships and for the range of 0-70 Nm/rad and 0-90 Nm/rad. Bolded values are significant, $p < 0.05$.

**Table S2** ANOVA P-values for 0-90 Nm/rad Range

| Name | Stance | Swing | Gait Cycle |
|---|---|---|---|
| Hip Flexion RMSE Mean | 0.1191 | 0.7098 | 0.1197 |
| Knee Angle RMSE Mean | **0.0159** | 0.2767 | **0.0316** |
| Ankle Angle RMSE Mean | **0.0007** | **0.0018** | **<0.0001** |
| Avg Joint RMSE Mean | **0.0023** | **0.0019** | **<0.0001** |
| Hip Flexion Work Net Mean | 0.8645 | 0.2453 | **0.0271** |
| Knee Angle Work Net Mean | 0.4990 | 0.7228 | 0.3752 |
| Ankle Angle Work Net Mean | **0.0091** | 0.2830 | **0.0194** |
| Total Work Net Mean | 0.0648 | 0.4781 | **0.0109** |
| Hip Flexion Work Pos Mean | 0.3628 | 0.1107 | **0.0092** |
| Knee Angle Work Pos Mean | 0.0988 | 0.5228 | 0.0849 |
| Ankle Angle Work Pos Mean | 0.2779 | 0.0804 | 0.4825 |
| Total Work Pos Mean | 0.5528 | **0.0135** | 0.2267 |
| Hip Flexion Work Neg Mean | 0.5454 | 0.3873 | **0.0190** |
| Knee Angle Work Neg Mean | **0.0003** | 0.6440 | 0.2507 |
| Ankle Angle Work Neg Mean | **0.0065** | **0.0302** | 0.0650 |
| Total Work Neg Mean | 0.0614 | 0.3792 | 0.5009 |
| Bicep Fem Total Mean | 0.4111 | 0.0729 | 0.2585 |
| Rec Fem Total Mean | 0.4921 | 0.2964 | 0.3772 |
| Ta Total Mean | **0.0185** | 0.7939 | 0.1938 |
| Soleus Total Mean | **<0.0001** | **0.0111** | **0.0001** |
| Gas Med Total Mean | 0.8084 | 0.7352 | 0.7809 |
| Bicep Fem Peak Mean | 0.2563 | 0.0775 | 0.1968 |
| Rec Fem Peak Mean | 0.6559 | 0.0893 | 0.3888 |
| Ta Peak Mean | **0.0003** | 0.8761 | **0.0005** |
| Soleus Peak Mean | **0.0021** | 0.2673 | **0.0033** |
| Gas Med Peak Mean | 0.7556 | 0.9776 | 0.7557 |
| Mean Activation Per Muscle Mean | 0.2275 | **0.0442** | 0.1260 |

ANOVA p-values for range of 0-90 Nm/rad stiffnesses. Bolded values are significant, $p < 0.05$.
